# Debottlenecking and reformulating feed media for improved CHO cell growth and titer by data-driven and model-guided analyses

**DOI:** 10.1101/2023.03.09.531884

**Authors:** Seo-Young Park, Dong-Hyuk Choi, Jinsung Song, Uiseon Park, Hyeran Cho, Bee Hak Hong, Yaron R. Silberberg, Dong-Yup Lee

**Author notes:** Correspondence to: D.-Y. Lee.

## Abstract

Designing and selecting cell culture media and feed are a key strategy to maximize culture performance in industrial biopharmaceutical processes. However, this is a major challenge for therapeutic proteins production since mammalian cells are very sensitive to their culture environment and require specific nutritional needs to grow and produce high-quality proteins such as antibodies. In this regard, in our previous study, we developed data-driven and in-silico model-guided systematic framework to investigate the effect of growth media on Chinese hamster ovary (CHO) cell culture performance, allowing us to design a new media formulation. To expand our exploration to feed, in this study, we evaluated two chemically defined feed media, A and B, in Ambr15 bioreactor runs using a monoclonal antibody-producing CHO K1 cell line. The feeds had a significant impact on cell growth, longevity, viability, and productivity and toxic metabolites production. Specifically, concentrated feed A was not sufficient to support prolonged cell culture and high titer compared to feed B. The framework systematically characterized the major metabolic bottlenecks in the TCA cycle and its related amino acid transferase reactions, and identified key design components, such as asparagine, aspartate, and glutamate, needed for highly productive cell cultures. From our results, we designed three new feeds by adjusting the levels of those amino acids and successfully validated their effectiveness in promoting cell growth and/or titer.

## 1 INTRODUCTION

Chinese hamster ovary (CHO) cells have been widely used as a mammalian workhorse for manufacturing recombinant therapeutic proteins (RTPs), including monoclonal antibodies (mAbs).^[1]^ For decades, substantial improvements have been made in their production by stable, robust, and high-producing cell lines generated through efficient vector design^[2,3]^ and cell screening and engineering systems,^[4,5]^ as well as improved culture processes^[6,7]^ and media development.^[8–10]^ Especially, designing culture media relevant to the specific cell line and culture process is a crucial step in early process development for achieving high RTP production with consistent and good quality during cell culture.^[11,12]^ Now, industrial-scale production of at least 2~8 g L^-1^ of RTP is expected, compared to protein titers of 0.2 g L^-1^ 20 years ago.^[13,14]^ In addition, media redesign or optimization benefits by fine-tuning certain media components to eliminate cellular bottleneck impacting the culture performance such as cell growth, viability, productivity, product quality and functionality.^[15,16]^ It is largely affected by the frequent accumulation of inhibitory metabolites or toxic by-products, e.g., lactate and ammonia, due to the inefficient CHO cell metabolism, which can be alleviated by adjusting certain media components as a source of nutrients and growth factors such as amino acids, glucose, vitamins, and trace metals.^[17–19]^

To provide an efficient manufacturing environment, each CHO cell culture medium typically contains 50-100 components, which are then tested and balanced with feed media using high-throughput experimental approaches.^[20]^ Traditional empirical methodologies, such as the one-factor-at-a-time (OFAT) approach^[21]^ and media blending^[22]^, can titrate single components but overlook the relationships among components. The design of experiments (DoE) methods, such as Plackett-Burman design and factorial design, are the conventional approach to streamline and improve screening designs by evaluating interactions of multiple media components and their impact on the culture performance, thereby reducing the overall experiment numbers required.^[23]^ However, these methods could be labor intensive, costly, and time-consuming, heavily relying on experimental data. Definitive-screening design (DSD) is an alternative one-step design method that requires a smaller number of experimental tests to identify main effects by two-factor interactions but requires more suboptimal media designs to be tested.^[24]^ Response surface methodology (RSM), such as Box-Behnken design^[25]^ or central composite design^[26]^, is utilized to investigate optimal concentrations and interactions of each component for well-informed or unknown processes, respectively. These DoE and RSM methods are useful for designing the most supportive media, but they are highly empirical and trial-and-error learning to explore multiple components with various concentrations, along with experimental replicates. In addition, these methods cannot be explained from a cellular metabolic perspective on culture performance improvement, so another media development strategy or analysis or verification experiment is needed to understand the complex interplay of media components in an intricate bioprocess. As the understanding of cell metabolism has grown, so has the development and formulation technology of culture media.^[27]^ Such disciplines as statistical analysis and genome-scale metabolic flux analysis have enabled efficient fine-tunning of nutritional requirements and harmonization of media and feeds. Spent cell culture media analysis and calculation of flux rate of nutrients and intermediates through complex cellular metabolic pathway are taken into account when the fine-tuning certain media components.^[28,29]^ Metabolic flux analysis has been applied to design the optimum composition and concentration of amino acids in CHO cell culture media, resulting in a dramatic increase in cell performance with 50% and 25% greater peak viable cell density and titer, respectively.^[30]^

More recently, it is desired to screen multiple media formulations and identify promising candidates followed by adjusting the component of culture media and process settings to achieve desired performance within a short timeline. Recent advancements in fully automated high-throughput screening systems, such as TubeSpin^®^ with Tecan EVO^®^ and Ambr^®^, have enabled rapid evaluation and design of complex media formulations.^[31,32]^ Despite the increased understanding and identification of the key components of CHO cell culture media through the aforementioned methods for media improvement, it still does not clearly establish a systematic approach that considers intricate interactions between multiple media components and moving targets in cellular metabolism during cultivation. In this regard, in our previous study, we developed data-driven and *in-silico* model-guided systematic framework to investigate the effect of growth media on CHO cell culture performance, allowing us to design the new media formulations for enhanced productivity.^[33,34]^

In fact, most of the biologics based on CHO cells are manufactured using fed-batch process. Since basal media may not contain all of the specific nutrients and growth factors required to support high levels of cell growth and productivity, feed media are typically supplemented to the basal media regularly during the culture process to provide additional nutrients and growth factors that are required for improving cell growth, productivity, and viability. The composition of the feed media is typically more complex than basal media and many include additional amino acids, vitamins, trace elements, and other supplements that are not present in the basal media. Additionally, the concentration of specific nutrients in the feed media may be higher than that in the basal media to ensure that the cells have access to the necessary nutrients over the culture process. By providing specific nutrients and growth factors at the right time and in the right concentrations, feed media can help to maximize the cell culture process and improve bioprocess performance. Designing and fine-tunning the feed medium is therefore a critical strategy for enhancing the yield of bioproducts while reducing the overall cost of production.

Thus, in this study, to investigate the impact of feed media on cell growth, productivity, and cellular metabolism during IgG-producing CHO K1 cell culture, we applied the modified version of a previously developed data-driven and *in silico* model-guided framework.^[35]^ This systematic media redesign framework involves four main hierarchical steps (**Figure 1**): culture data collections, multivariate statistical analysis, *in silico* flux balance analysis (FBA) with CHO-specific genome-scale metabolic model, and targeted identification of critical nutrients based on knowledge to identify and remove bottlenecks that limit cell growth or productivity. By systematically identifying key media components and addressing recurrent metabolic bottlenecks, this framework can be used routinely applied for media redesign and reformulation in biologics process development. Ultimately, it could be an effective approach for promoting consistent cell growth and productivity in the biopharmaceutical industry.

**FIGURE 1.**
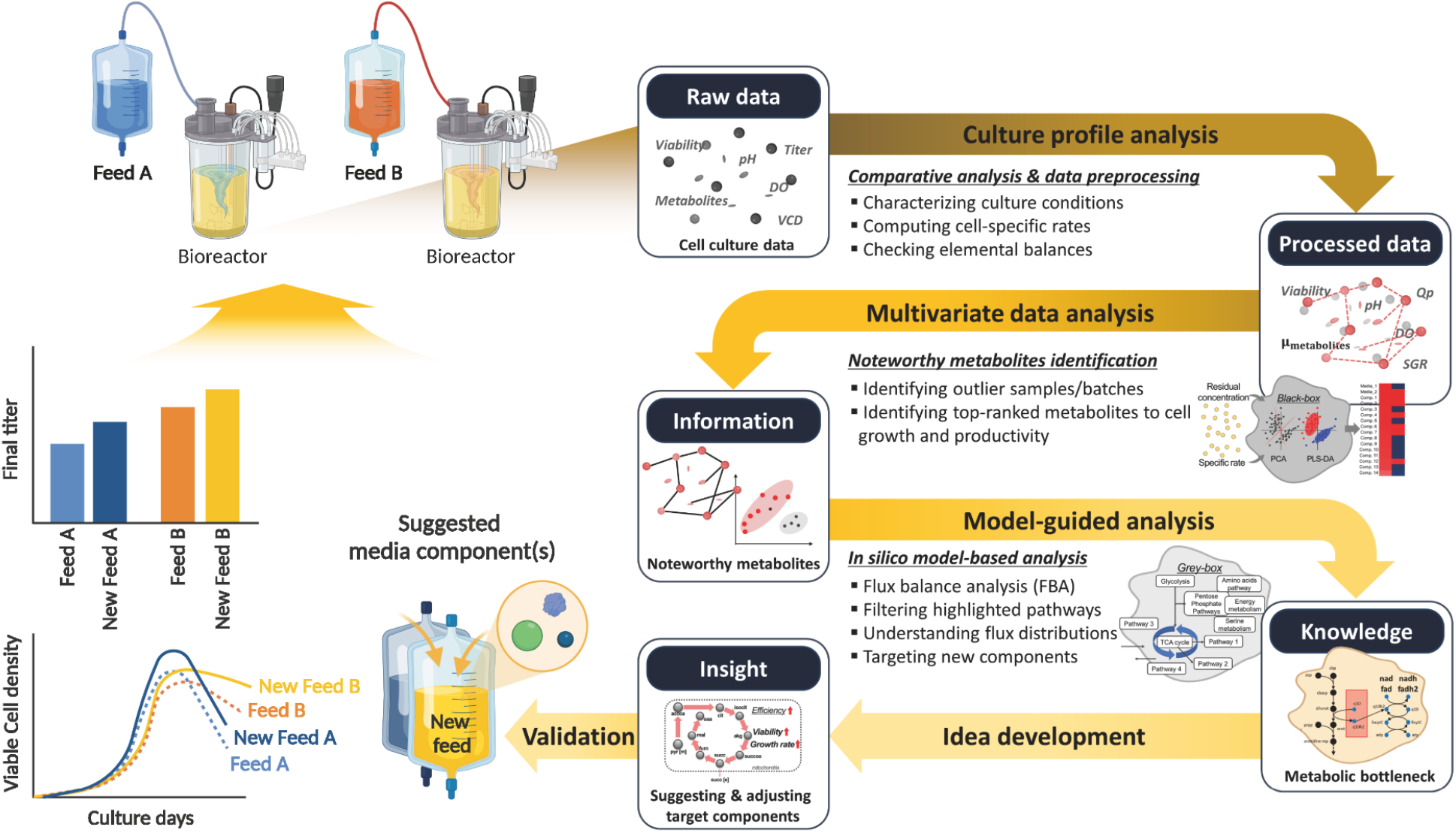
Scheme of a systematic framework for data-driven and model-guided analysis for elucidating media performance and development. Cell culture and metabolite profiles are collected and preprocessed. MVDA and FBA on GEM analyses are then utilized to identify key design components that improve culture performance.

## 2 MATERIALS AND METHODS

### 2.1 CHO cell culture conditions and profiling

A proprietary recombinant CHO-K1 cell line was used to produce an IgG1 antibody. For the seed trains, one frozen vial from a working cell bank was thawed in a proprietary chemically-defined CELLiST™ (Ajinomoto Genexine, Co., Ltd.) basal medium and cultured for two consecutive passages. This was followed by three passages in a 125 ml Erlenmeyer shake flask (Corning Life Sciences, New York, USA) with 30 mL working volume at a seeding density of 3 x 10^5^ cells mL^-1^. The flasks were placed on an orbital shaker (22 mm diameter; N-biotek, Republic of Korea) in an incubator at 37°C with agitation rate 115 rpm and humidified atmosphere of 5% CO_2_ before the production run. The culture was supplemented with 2 mM of L-glutamine (Thermo Fisher, Waltham, MA, USA), 30 ug mL^-1^ of puromycin (Thermo Fisher, Waltham, MA, USA), and 5 mg L^-1^ of insulin (Sigma-Aldrich, St. Louis, MO, USA).

The 15 mL fully automated small-scale ambr15^®^ bioreactor (Sartorius, Göttingen, Germany) was used for 14 days production run with 12 mL working volume under two proprietary different feed media supplements, referred to as feed A and B. Both basal and feed media used were from Ajinomoto Genexine’s ‘CELLiST’ cell culture media panel. The fed-batch culture with feed A and B were conducted in duplicate and triplicate, respectively, and the cultures were incubated at 3 x 10^5^ cells mL^-1^ with feed was supplemented every two days, starting from day 4, at 4% (v/v). A bolus glucose was added up to 6 g L^-1^ for day 4-8 or 7 g L^-1^ for day 9-12 when the residual glucose concentration was below 4 g L^-1^. All physical process parameters were monitored and adjusted simultaneously in real-time via ambr^®^ runtime software as 37°C, stirrer speed of 1200 rpm, pH 7.0±0.05 and DO 50%. The pH was controlled by the addition of CO_2_ to reduce the pH or addition of 7.5% (w/v) sodium carbonate to increase the pH. Additionally, 1% (v/v) Antifoam C Emulsion (Signa-Aldrich, St. Louis, MO, USA) was added depending on the foaming level to avoid foam overflow.

The viable cell density (VCD) and viability were measured using a Cedex HiRes Analyzer (Roche, Basel, Switzerland) from daily samples. Samples collected on days 0, 2, 4, 6, 7, 8, 10, 12 and 14 were analyzed using a Cedex Bio Analyzer (Roche, Basel, Switzerland) to monitor glucose, glutamine, glutamate, lactate, ammonia, and IgG concentration. The concentration of amino acids in the supernatant were determined using the AccQ-Tag Ultra Derivatization kit (Waters, Milford, MA, USA) as per the manufacturer’s instructions. The derivatized sample was then analyzed using an ACQUITY H-class ultra-high-performance liquid chromatography system (Waters, Milford, USA) with a 1.7 μm 2.1 x 100 mm AccQ-Tag ultra C18 column (Waters, Milford, MA, USA). The amino acids were separated on the column at 43°C with a gradient method. Mobile phase A consisted of 100% AccQ-Tag Ultra Eluent A, mobile phase B was a 0.1% (v/v) AccQ-Tag Ultra Eluent B in water, mobile phase C was 100% UPLC grade water, and mobile phase D was 100% AccQ-Tag Ultra Eluent B. The flow rate was set to 0.7 mL min^-1^, the gradients of mobile solutions are described in **Supporting information Table S1**. The separated amino acids were detected using an Acquity UPLC^®^ Tunable UV (TUV) detector at 260 nm, and the concentration of amino acids were quantified using the Empower software (Waters, Milford, MA, USA).

### 2.2 Collected data quality check and multivariate data analysis

To determine the media components that contribute to the difference in culture performance over time, we calculated the specific growth rate (SGR, mmol gDCW^-1^ hr^-1^) and specific productivity (Qp, pmol cell^-1^ day^-1^) using the method described in the previous study.^[36]^ The concentrations of metabolites such as glucose, lactate, ammonia, amino acids were also converted into cell-specific consumption or secretion rates between each time point. The preprocessed data were assorted by time points (day 0, 2, 4, 6, 7, 8, 10, 12, and 14) and phases (days 0-2, 2-4, 4-6, 6-7, 7-8, 8-10, 10-12, and 12-14) for carbon elemental balancing and then analyzed using a series of multivariate data analysis (MVDA) technologies with SIMCA^®^ software (vers.16, Sartorius-Stedim, Umea, Sweden). Principal component analysis (PCA) was performed to identify any outlier samples and to provide an overview of the relationship between two feed media conditions. To specifically examine the effect of the feeds on SGR, four phases (days 4-6, 6-7, 7-8, and 8-10), which displayed significantly different SGR after feeding on day 4, were selected for further analysis. Next, we performed a batch evolution model (BEM)-based orthogonal partial least square (OPLS) analysis to evaluate the effect of specific metabolites rates (input variables, X) on SGR (output variable, Y) at individual time points across each reactor condition, allowing for the identification of underlying time-based trends. Variable influence on projection (VIP) scores were used to quantify the importance of X-variables on the Y variable. Metabolites with VIP scores greater than 1.0 for the four selected phases and high VIP scores in all three phases, where media components differed by more than two-fold between feed A and B, were filtered and selected for further analysis. These metabolites were analyzed in-depth using a genome-level mechanistic model.

### 2.3 *In silico* ecFBA with CHO-specific GEM

To investigate how differences in feed media affect metabolic flux changes and identify which cellular metabolism is crucial for these differences and can be manipulated to enhance cell growth, we utilized the latest CHO genome-scale metabolic network model (GEM), *i*CHO2291, with enzyme-capacity constrained FBA (ecFBA) by incorporating both measured cell-specific rates (glucose, lactate, ammonia, amino acids, productivity) and enzyme kinetic parameters (*k*_cat_).^[37]^ We performed ecFBA using the constraint-based reconstruction and analysis (COBRA) Toolbox vers.3.0,^[38]^ implemented in MATLAB (R2020a, MathWorks) with Gurobi vers.9.1.1 (http://www.gurobi.com) as the solver for linear programming problems. Specifically, we studied the after feeding- and exponential-phases on days 6-7, which exhibited the most differentiated SGR with similar cell-specific productivity. The biomass objective function was maximized in overall simulations and the constant rate of the measured productivity was set as a constraint. To eliminate the mathematical infeasible solution space and account for experimental perturbation, both upper (+10%) and lower (−10%) bounds were relaxed by ±10% with respect to the uptake rates of nutrients.

To determine which metabolic networks to investigate, we analyzed and visualized the flux ratio of individual reactions within the CHO genome-scale metabolic network as a map using Cytoscape (vers.3.9.1, https://cytoscape.org)^[39]^ and calculated the fold change (*FC*) values to determine the flux differences between the two feed conditions using equation (1). We neglected the flux value of reactions below 0.0001 as shown in equation (2). The top 50 most distinctive metabolic reactions among 4,334 global reactions with non-zero fluxes were selected based on equation (3), with *FC* values above 1.2.

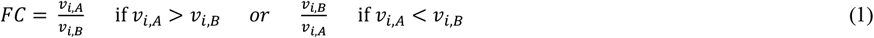

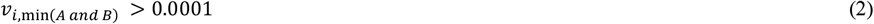

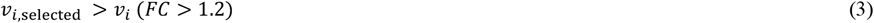

Where *v_i,A or B_* is the flux of the reaction *i* in feed A or B condition. *v_i,min_* is the minimum flux value under A and B conditions.

### 2.4 Validation experiment

To validate our systematic approach, we reformulated three new feed media by adjusting the concentration of specific amino acids that had the strongest correlation with the bottlenecked metabolism that could potentially influence cell growth. As a proof of concept, each feed medium was formulated by changing the level of each target amino acid, namely asparagine, aspartate, and glutamate. A 50% increase in the level of glutamate was added to the original feed A for duplicate bioreactor runs, while a 100% elevation in the level of asparagine and aspartate were added to the original feed B for triplicate runs. The same ambr15^®^ bioreactor system and operating conditions were applied for the experimental validation runs.

## 3 RESULTS

### 3.1 Systematic framework for characterizing media-dependent culture and identifying adjustable media component targets

In this study, in order to identify key component targets of feed media for the enhanced cell culture performance or reduced toxic effect of waste byproducts, we applied data-driven and *in silico* model-guided systematic framework which has been successfully used to explore the effect of cell-line specific basal media on cellular metabolism and design a new media formulation for improving cell growth.^[34]^ We have simplified this framework by utilizing only the common daily measurement and amino acids concentration data in biologics upstream process, which consists of four hierarchical steps as shown in **Figure 1**. First, general cell culture profiles such as VCD, viability, titer, and common metabolite concentrations including amino acids are collected from CHO cell cultures in two different feed media conditions. The profiled experimental data is examined to gain insight into the quality of the dataset and then converted to cell-specific consumption or secretion data, which quantifies the rates of growth, target RTP production, and metabolite utilization or formation per cell and culture time unit. The elemental balance of main inputs (nutrients) and outputs (products) is checked and corrected. Next, the processed data undergoes MVDA to find the correlations between metabolic exchange rates and SGR. PCA is used to provide an overview of the differences in culture between the two feed conditions, and then BEM-based OPLS analysis is applied to further analyze the time-dependent cellular activity and metabolic behavior that may explain the effect of feed media on cell growth. The most influential components are identified via an OPLS-based VIP plot. Model-guided analysis is then carried out using ecFBA to delineate the metabolic state differences (i.e., flux distributions) that may be caused by specific metabolites and blocked reactions in the metabolic network. The metabolites and their relevant critical metabolic pathways can be identified, which make major difference in the culture and metabolic behavior in feed media dependent conditions. Finally, to eliminate cellular bottlenecks and support the cell’s metabolic needs, target metabolites that are positively associated with growth and productivity can be suggested for adjusting the concentration of specific nutrients or changing the medium composition as a new media formulation.

### 3.2 Cells cultured under a less concentrated feed media condition show better IgG production

To examine the impact of the feed medium on cellular growth and specific productivity, IgG1-producing CHO-K1 cell line was cultured in strictly controlled ambr15^®^ bioreactors for 14 days using the same basal medium and two chemically-defined proprietary feeds A and B, which were derived from the manufacturer’s media development procedure (Ajinomoto Genexine, CELLiST product). The culture profiles were collected, preprocessed, and summarized in **Figure 2**. Until day 4, the culture profiles under both feed conditions were very similar. However, after feeding on day 4, significant differences emerged between two conditions in all profiles. The cells cultured in feed A grew faster and to a higher density, with a higher SGR until day 8, and reached peak VCD of 35.9 x10^6^ cell mL^-1^ on day 9 (8% difference), one day earlier than feed B. After reaching peak VCD, the cells declined rapidly with cell viability dropping dramatically to 49.78% on the last day of culture in feed A. In contrast, the VCD of feed B decreased slowly and maintained a clear plateau phase of about four days (i.e., longer longevity), with cell viability remaining above 88% throughout the entire culture duration. The feed media also impacted the titer and specific productivity (Qp) with differences observed starting from day 10. Feed B showed higher titer (690 *vs*. 777 mg L^-1^ and 752 *vs*. 791 mg L^-1^ on days 12 and 14, respectively) and higher Qp, which was slightly lower until day 8 but substantially higher (15% difference) on day 10.

**FIGURE 2.**
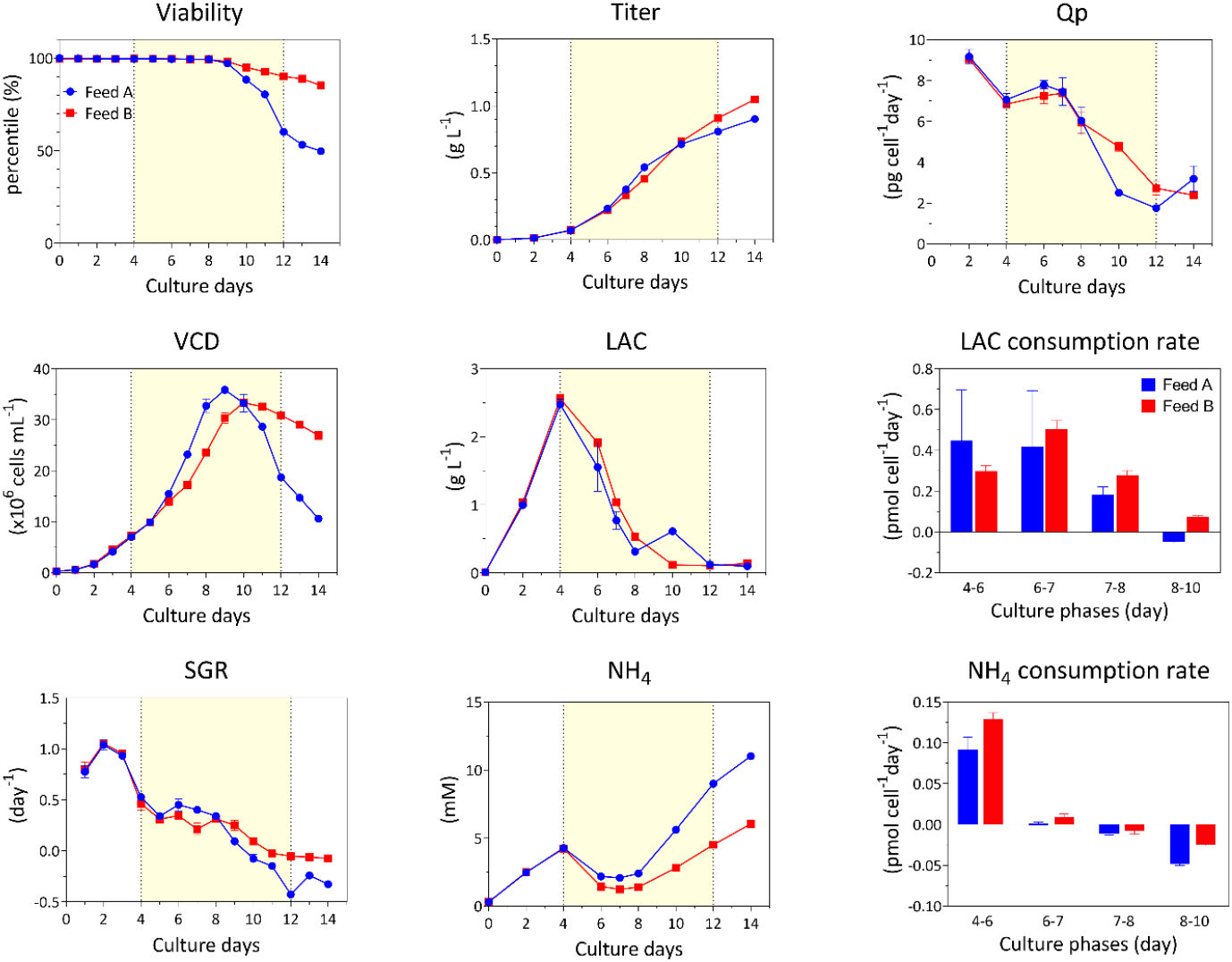
CHO cell culture profiles in two different feed media. Effect of feed media on viable cell density (VCD), viability, specific growth rate (SRG), titer, specific productivity (Qp), lactate and ammonia (NH4) and their specific rates over 14 days CHO cell cultures. Feed media added from day 4 to 12 (yellow shade). Error bars show standard deviation across duplicate or triplicate cultures.

The metabolites profiles in the two feed conditions showed considerable differences (**Figure 2, Supporting information Figure S1**). In the higher titer condition B, more lactate was accumulated and subsequently consumed after feeding started on day 4. The specific consumption rate was lower on days 4-6 compared to feed A, and vice versa behavior was observed between days 6-14, implying that high lactate consumption in the late phase of culture can contribute to increase production.^[40]^ Interestingly, the ammonia concentration increased and slightly decreased on day 4, and then continued to increase throughout the cultivation in all conditions. However, in the higher titer condition B, a higher ammonia consumption rate was observed between days 4-6, and much lower secretion rate was observed until the end of culture, while much higher production was observed in the lower titer condition A. Thus, these toxic byproduct levels in each feed condition likely contribute to the differentiation in culture performance.

### 3.3 Multivariate data analysis identifies differences in nutrients usage after feeding phase

The overall difference in cell culture profiles was confirmed to be evident even when using different feed media. Compared to feed A media, the use of feed B media resulted in superior cell growth, longevity, and productivity (Qp). To identify which media component induced the difference in culture performance over time, nutrients concentrations were compared to the composition of the feed medium used. Most nutrients were more concentrated (with an average 38% higher concentration) in feed A medium, while alanine (100%), glutamate (53%), and isoleucine (38%) concentrations were higher in feed B (**Figure 3A**). Asparagine, aspartate, and methionine were more than 3.1-, 2.5-, and 4.3-folds higher in feed A, respectively. Aspartate, glutamate, and tryptophan were quickly consumed and almost depleted on day 8 or 10 in the feed A condition (**Supporting information Figure S1**). Asparagine, cysteine, and leucine were consumed on day 4, 6, 10, respectively, in both feed media. Only methionine (77% higher in feed A medium) was gradually utilized and exhausted on day 10 in the feed B condition.

**FIGURE 3.**
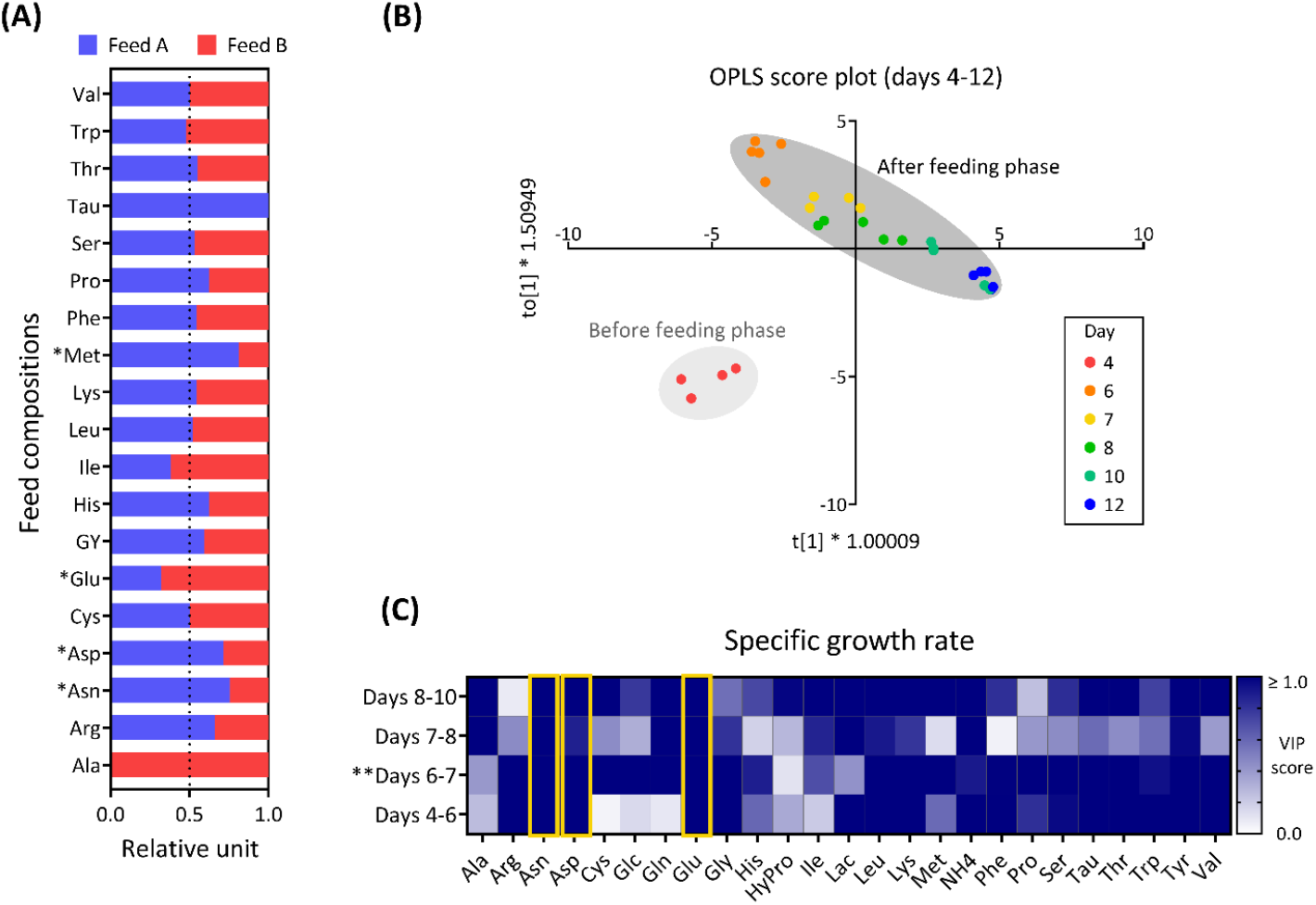
**(A)** Comparison of components in two different feed media A and B. **(B)** Score plot of metabolite concentration was carried out via BEM-based OPLS analysis to identify the effect of specific rates of amino acids, glucose, lactate, and ammonia on SGR throughout the course of the culture. **(C)** Heatmap of VIP score for correlation individual metabolite exchanges with SGR in different culture phases.

To investigate how these differences in measured metabolites are related to cell growth and titer during the culture, BEM-based OPLS was performed using cell-specific consumption or secretion rates with aligned time points across each reactor condition (two batches for each feed condition). The score plot in **Figure 3B** shows the multiple time points within a batch across different feed media. This confirms that bioreactors operated under the same conditions perform similarly. The duplicate bioreactor conditions tend to cluster, and the maturity of cultures at each time point is represented by different colors. Since it clearly shows two clusters, i.e., before and after feeding phases based on day 4, we focused on analyzing the four phases in after feeding phase, i.e., days 4-6, 6-7, 7-8, and 8-10 to determine which and how the various components contribute to the separation of the clusters and help to improve the specific growth. Key identified metabolites from the VIP score (threshold ≥1) plot are asparagine and glutamate in all four phases as well as aspartate in all three phases (**Figure 3C**). Therefore, to promote the specific growth, a feed should be designed with higher asparagine, aspartate and/or glutamate concentration depending on feed type. These amino acids are to be the candidates for the most significantly growth-affecting components being analyzed for further genome-level mechanistic model analysis.

### 3.4 ecFBA with CHO GEM represents feed-dependent metabolic flux distribution in late glycolysis to TCA cycle and glutaminolysis reactions

The ecFBA of *i*CHO 2291 was conducted to mechanistically investigate how feed media differences drive metabolic flux changes and to identify which cellular metabolism is the major difference that could be manipulated to enhance cell growth. Specific uptake and section rate of metabolites during the exponential and after feeding phases were considered, and then a global and unbiased assessment of flux distributions in the CHO metabolic network was performed. **Figure 4** illustrates the most prominent metabolic pathways distinguishing feed A from feed B, identified through a systematic analysis. A higher fold change of pathways in either A (blue) or in B (red) indicates that the amino acid composition in each feed is driving different metabolic traits in cells through corresponding reactions and pathways. This feed-dependent metabolic response is observed in several CHO metabolic pathways during after feeding phase, especially the early exponential phase (days 6-7). A total of 50 reactions and their associated pathways, mainly in central carbon and amino acids metabolism, were identified with fold-changes greater than 1.2. To investigate how these metabolic changes occur in central carbon and amino acids metabolism between the two feed conditions, we predicted intracellular fluxes. The resulting analysis showed that these fluxes were differentially distributed in the two feed conditions, as shown in **Figure 5A**.

**FIGURE 4.**
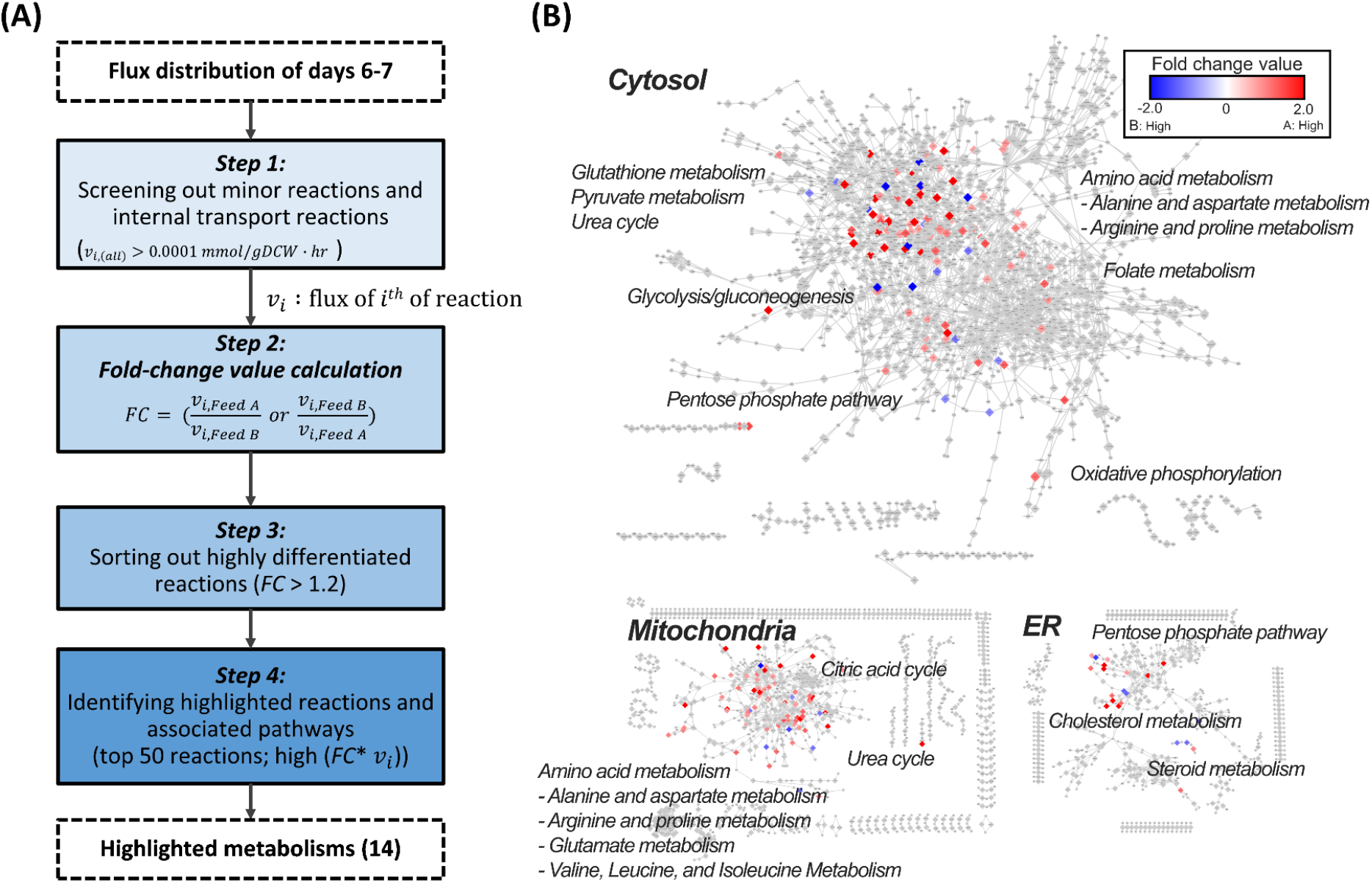
**(A)** Steps for isolating the differentiated metabolic pathways between two feed coditions during day 6-7. **(B)** The resulting metabolic network in cytosolic, endoplasmic reticulum (ER) and mitocondria was visualized using Cytoscape. Colored dots indicate the significance of fold change of flux values in particular subsystem.

**FIGURE 5.**
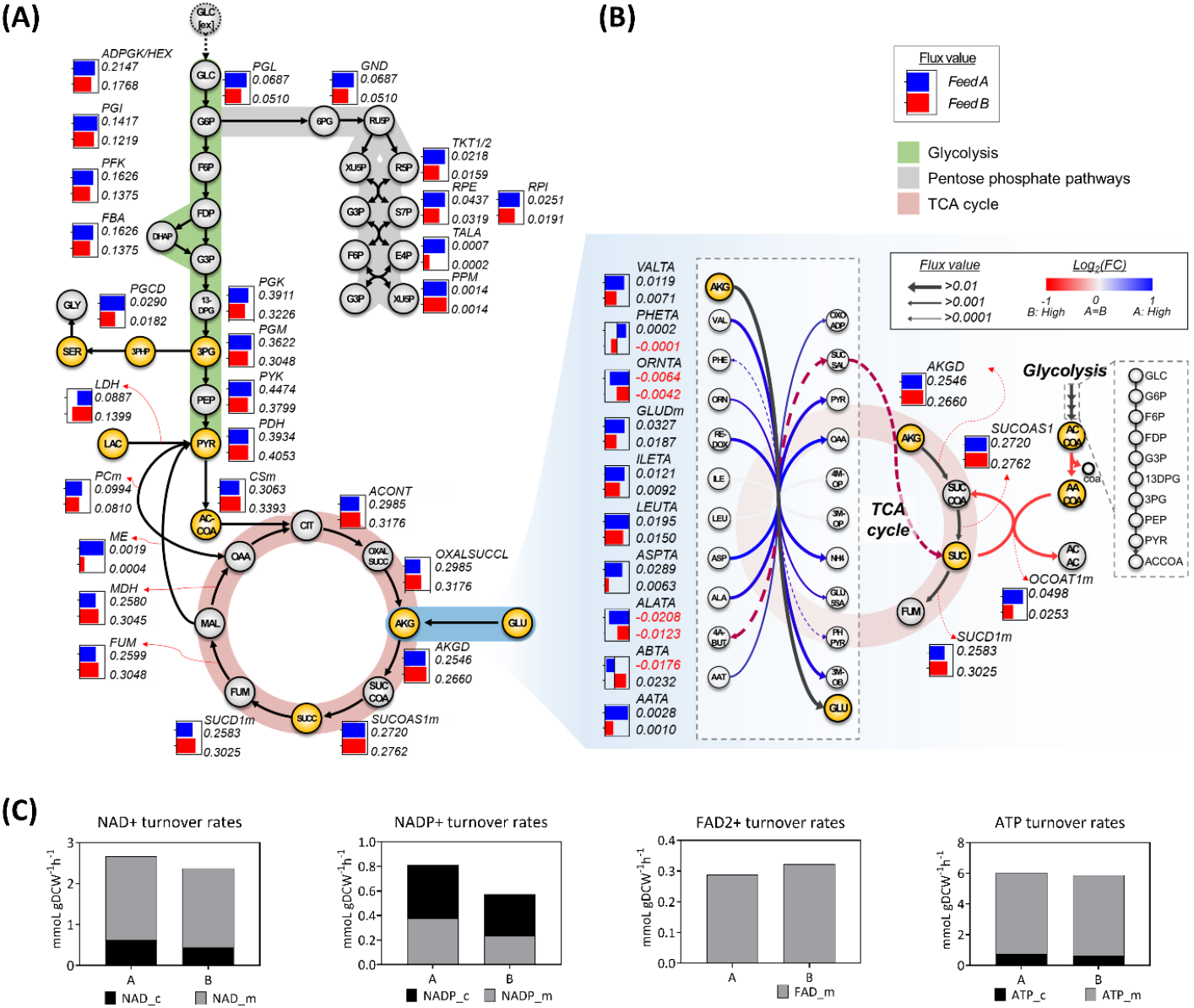
**(A)** Overall metabolic flux distribution in central carbon metabolism and **(B)** alpha-ketoglutarate and glutamate pathways, and **(C)** turnover rates of redox cofactors such as NAD, NADP, FAD, and ATP respectively in two feed conditions.

Distinct flux changes (*FC* >50%) were observed in specific metabolic pathways including glycolysis, pentose phosphate pathway (PPP), serine pathway, lactate pathway, tricarboxylic acid (TCA) cycle and its intermediate pathways, in response to the different feed conditions. In the feed A condition, which supported higher cell growth, there was a higher glycolytic flux until pyruvate, with a significant bypass through PPP. Additionally, the A condition exhibited intensified glycolysis-diverting flux (1.6-fold higher flux rate) in phosphoglycerate dehydrogenase (PGCD), which consumes the glycolytic intermediate 3-phosphoglycerate (3pg) to supply serine and generate NADPH through three-step enzymatic reactions. The flux of pyruvate kinase (PYK) in glycolysis was also high in the A condition, which regulates glycolytic flux and produces ATP. This suggests that cells in the A condition may use glycolysis to enhance cell proliferation and fast growth via serine biosynthesis, but the cultivation showed decline in culture longevity, cell viability, and productivity (as seen in **Figure 2**). This might be due to a reduced supply of caron source to the TCA cycle and a lower utilization of the TCA cycle. In contrast, the B condition, which supported higher production, exhibited different flux in the last step of glycolysis, involving pyruvate metabolism with higher flux rates of pyruvate dehydrogenase (PDH) and lactate dehydrogenase (LDH), leading to a substantial influx to the TCA cycle. Additionally, there was a higher lactate consumption rate in B condition, as indicated by the lactate profile.

Among the series of enzymatic reactions in the TCA cycle, the metabolic behaviors of glutamate-alpha-ketoglutarate (AKG) metabolism showed a distinct pattern between the two conditions (**Figure 5B**). AKG is essential for the synthesis of glutamate, which is important for balancing highly consumed metabolites through various amino acid transferases. In feed A condition, a reversible reaction was observed where glutamate converted to AKG and ammonia, while glutamate was supplied from abundant TCA intermediates in feed B condition. It is important to note that feed A had 53% lower concentration and uptakes of glutamate compared to feed B. Glutaminolysis, which is composed of amino acid transferases, such as aspartate transaminase (ASPTA), aminobutyrate transaminase (ABTA), and mitochondrion glutamate dehydrogenase (GLUDm), reversibly convert glutamate to AKG, resulting in high-energy redox cofactors such as NADP more regenerated in condition A (**Figure 5C**). Among these reactions, aspartate was the most significant contributor to the generation of AKG, especially in the lower production condition A (ASPTA 0.0289 *vs*. 0.0063 mmol gDCW^-1^. Higher uptake of asparagine and aspartate in condition A boosts conversion AKG to glutamate, allowing cells in A condition to be supplied with the necessary glutamate due to its deficiency in media. TCA intermediates are also coupled with aceto-acetyl coenzyme A (aacoa) and acetoacetate (acac) in glycolysis via 3-oxoacid CoA-transferase (OCOAT1m). Interestingly, marginal flux differences of succinyl-CoA ligase (GDP-forming) (SUCOAS1m) became enlarged due to the high ABTA flux, which is a part of the glutamate-AKG conversion. This phenomenon was due to the complex interconnected network in the glutamate-AKG conversion, possibly induced by the deficient asparagine, aspartate, and glutamate.^[12]^ Therefore, we further investigated the glutamate-related metabolic reactions, where these amino acids are interconnected in the metabolic pathways (**Figure 6**).

**FIGURE 6.**
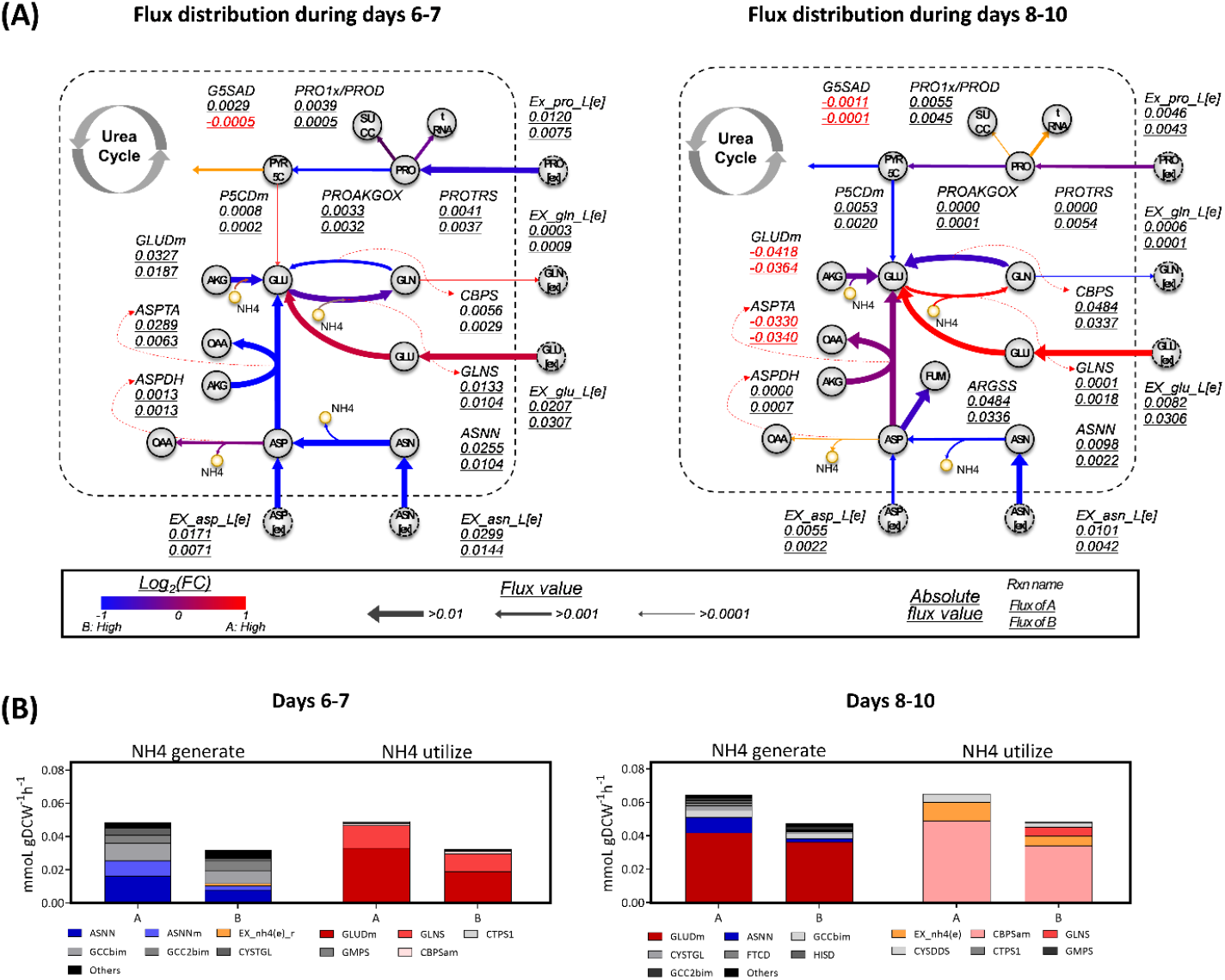
Asparagine, aspartate, and glutamate metabolism. **(A)** Flux distributions of alpha-ketoglutarate and glutamate relatived reactions during two different culture phases (days 6-7 *vs*. days 8-10) and their **(B)** overall ammonia turnover rates.

Ammonium (NH_4_) accumulation and glutamate depletion were observed in the late stage of the exponential phase in condition A, where the VCD suddenly drops. We estimated the turnover rates of NH_4_, which indirectly indicates the metabolic pool of NH_4_ with the flux-sum approach (**Figure 6A**). During days 6-7, GLUDm contributed to NH_4_ utilization, while asparaginase participated in NH_4_ generation (**Figure 6B**). Most of the asparagine- and aspartate-relevant reactions were enriched in condition A, possibly due to higher concentration of aspartate and asparagine in feed A compared to feed B. We also investigated flux distribution from days 8-10 to estimate the role of glutamate and identify the differentiated flux distribution of NH_4_. The direction of glutamate dehydrogenase (GLUD) changed from NH_4_ utilization to generation, causing subsequent NH_4_ accumulation in condition A. Therefore, we hypothesize that the depletion of glutamate in condition A causes intracellular synthesis of glutamate on days 6-7, subsequently inducing NH_4_ accumulation by changing the role of GLUDm. Hence, we suggest increasing glutamate concentration to sustain a reliable NH_4_ level in condition A and increasing depleted amino acids such as asparagine or aspartate to achieve more cell proliferation in condition B. To validate this hypothesis, we performed validation bioreactor runs by adding single amino acids to each feed: glutamate in feed A and asparagine and aspartate in feed B (**Figure 7**). The addition of glutamate to feed A significantly promoted cell growth (9% peak VCD), viability (average 32%), and final titer (19%) while reducing NH_4_ accumulation. The supplementation of asparagine to feed B led to a 7% increase in peak VCD and a 5% increase in final titer, while maintaining comparable viability. Similarly, in the case of addition aspartate to feed B, a slight increment of 2% was observed in both peak VCD and final titer compared to the previous feed B, with no significant impact on cell viability.

**FIGURE 7.**
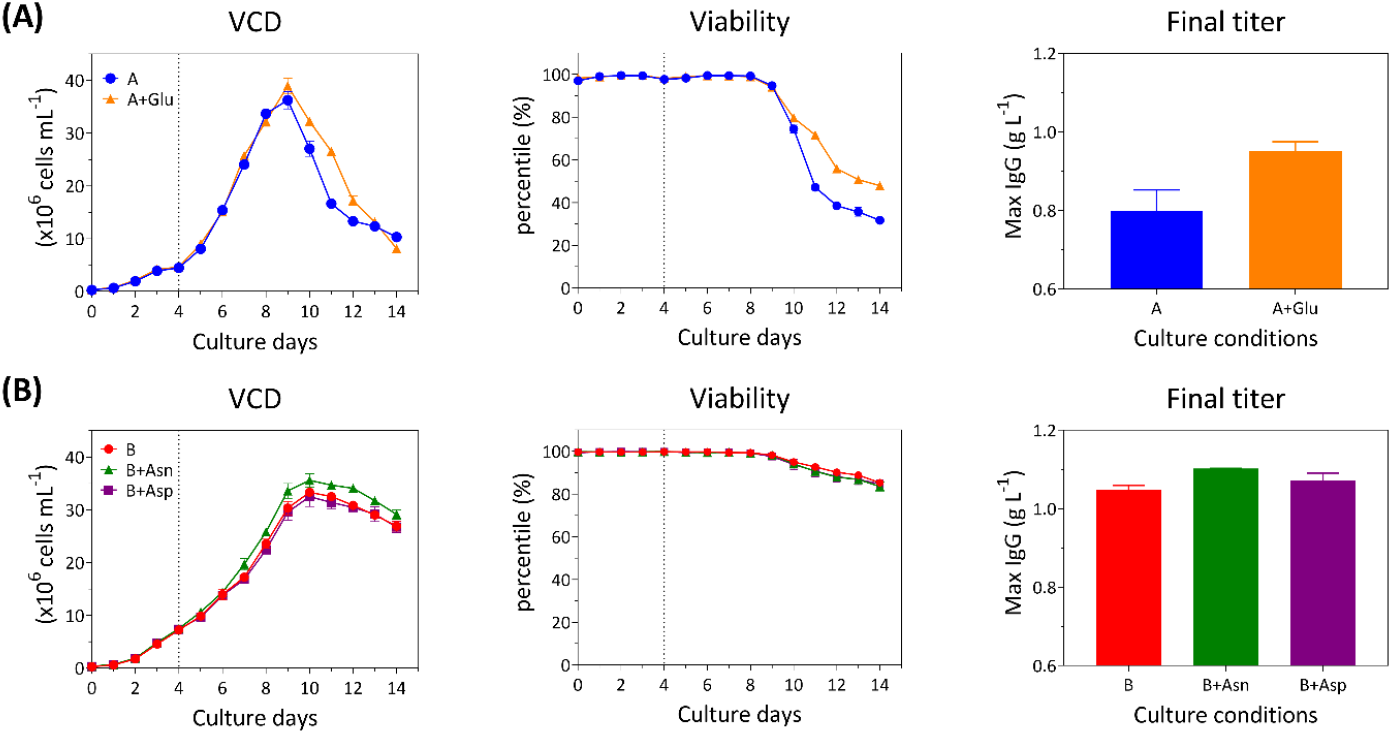
The validation experiment demonstrated that the identified specific amino acids for **(A)** feed medium A with supplemented glutamate (Glu) and **(B)** feed medium B with asparagin (Asn) and aspartate (Asp) were able to improve the culture performance. Error bars show standard deviation across duplicate or triplicate cultures.

## 4 CONCLUSIONS

In this study, we successfully applied our previously developed data-driven and *in silico* model-guided systematic framework to investigate the impact of feed media on culture behavior and identify metabolite targets for improving feed media design. This study builds upon our earlier work, which focused on the effect of basal (growth) media on cell culture performance. We addressed the limitations of two different feed media in the CHO cell culture process by adjusting the concentration of specific nutrients in the feed to better support the cells’ metabolic needs throughout cultivation. Initially, CHO cells were cultured in two different feeds and the culture profiles were collected. We then used multivariate statistical analysis to identify noticeable components and metabolite exchanges that are correlated with culture performance (specific growth or productivity). Following this, FBA of CHO GEM was conducted to elucidate the metabolic mechanisms involved and to find blocked reactions in the metabolic network. Finally, the validation bioreactor runs confirmed the key metabolites and involved reactions identified by data-driven and model-guided analyses for debottlenecking and reformulating feed media to enhance cell growth, longevity, and titer.

## 5 CONFLICT OF INTEREST

The authors declare no conflict of interest.

## 7 FIGURE LEGENDS

**FIGURE S1.**
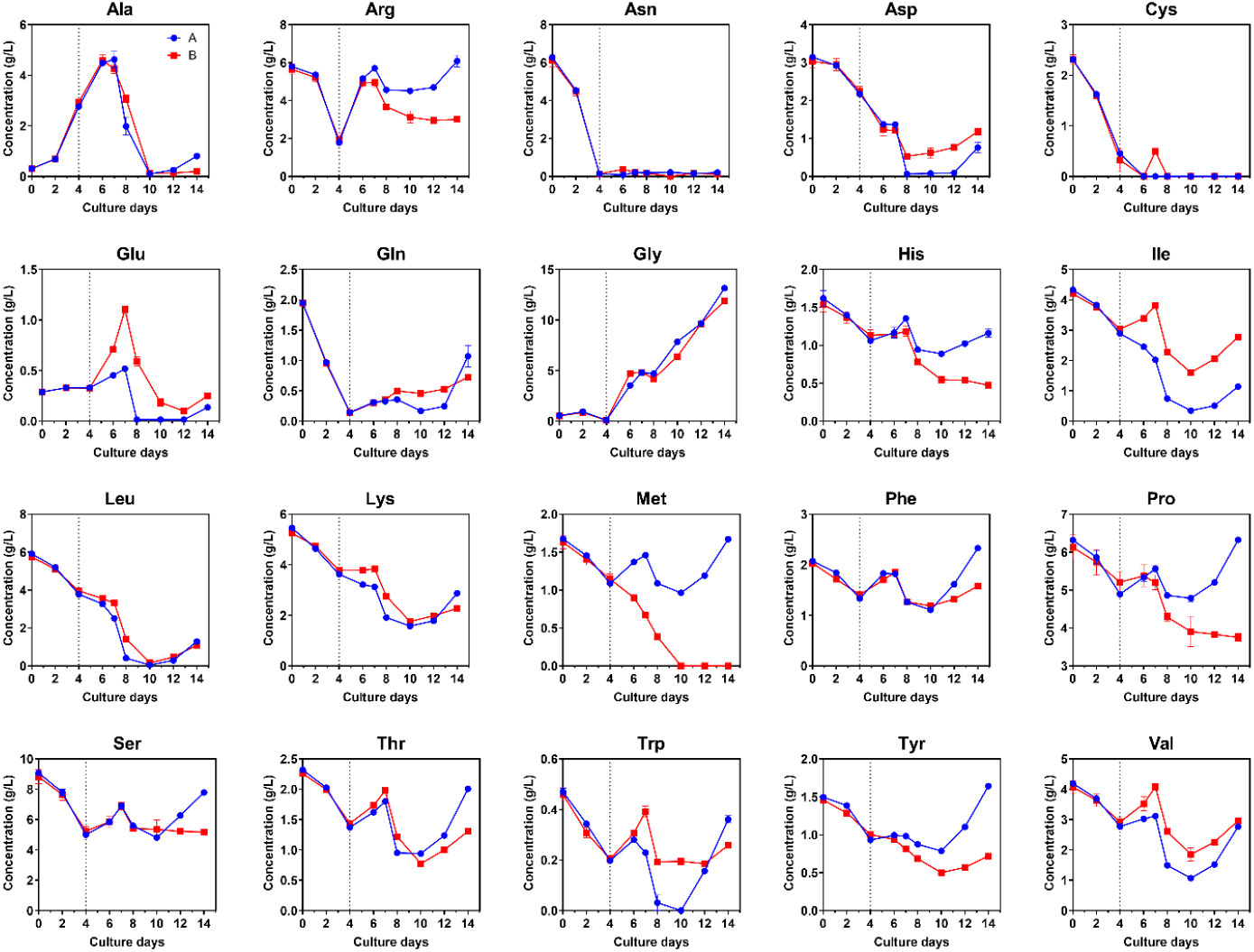
Residual amino acids concentration profiles over 14 days CHO cell cultures.

**Supplementary Table S1.** Gradient profile applied in the developed UPLC method

| Time (min) | Mobile A (%) | Mobile B (%) | Mobile C (%) | Mobile D (%) | Curve |
| --- | --- | --- | --- | --- | --- |
| 0.00 | 10.0 | 0.0 | 90.0 | 0.0 | N/A |
| 0.29 | 9.9 | 0.0 | 90.1 | 0.0 | 11 |
| 5.49 | 9.0 | 80.0 | 11.0 | 0.0 | 7 |
| 7.10 | 8.0 | 15.6 | 57.9 | 18.5 | 6 |
| 7.3 | 8.0 | 15.6 | 57.9 | 18.5 | 6 |
| 7.69 | 7.8 | 0.0 | 70.9 | 21.3 | 6 |
| 7.99 | 4.0 | 0.0 | 36.3 | 59.7 | 6 |
| 8.59 | 4.0 | 0.0 | 36.3 | 59.7 | 6 |
| 8.68 | 10.0 | 0.0 | 90.0 | 0.0 | 6 |
| 10.20 | 10.0 | 0.0 | 90.0 | 0.0 | 6 |

## Notes

### Competing Interest Statement

The authors have declared no competing interest.

